# Dissection of figured wood trait in curly birch (*Betula pendula var. carelica*) using high-throughput genotyping

**DOI:** 10.1101/2023.11.07.566062

**Authors:** Rim Gubaev, Dmitry Karzhaev, Elizaveta Grigoreva, Kirill Lytkin, Elizaveta Safronycheva, Vladimir Volkov, Veronika Nesterchuk, Lidiya Vetchinnikova, Anatoly Zhigunov, Elena Potokina

**Author notes:** Gregor Mendel Institute of Molecular Plant Biology, Vienna, Austria (Elizaveta Grigoreva).

## Abstract

Curly (Karelian) birch is a special variety of *Betula pendula* distributed in the northwestern part of Europe. Karelian birch is well-known for its valuable figured curly wood also known as “wooden marble”. The genetic basis underlying curly wood formation has been debated since last century, however, there was no data about loci responsible for the curly wood trait. In the present study, we analyzed two full-sibs populations derived from experimental crosses of curly birches and segregating for the trait. RADseq genotyping was applied to reveal how many loci are involved in ‘curliness’ formation and to search for genetic variants associated with this trait. One single interval on chromosome 10 was detected containing possible candidate genes. InDel marker BpCW1 was suggested for the first time for marker-assisted selection of trees with curly wood at their earliest stages of development.

## 1 Introduction

Curly (Karelian) birch (*Betula pendula* Roth var. *carelica* (Mercklin) Hämet-Ahti) is a special variety of silver birch, which forms small populations scattered within the north-western part of *B. pendula* distribution: Finland, Russia (Karelia), Baltic countries, Belarus, Poland, southern Sweden and Norway ^1,2^. The wood fibers of the Karelian birch are not directed strictly vertically, but at different angles, which leads to the formation of wood “curliness” underlying the special ‘curly’ phenotype. The figured ‘curly’ wood of the Karelian birch is used for manufacturing highly decorative furniture and architectural panels, and it is sometimes called ‘wooden marble from Finland’ ^3^. Because of its exotic nature, figured wood is paid by weight (3-5 € kg^-1^), rather than by volume like the wood of other trees ^4^. Due to the high commercial value, interest in the propagation of curly birch on plantations has increased significantly since the 1980s. In Russia, the total area of forest plantations of Karelian birch by 1986 reached 5500 ha ^5^. Unfortunately, in the 90s, due to illegal logging, about 1.5 thousand curly birches were cut down on the territory of Karelia. In Finland, though, 6500 ha of curly birch stands have been established, and they are currently reaching their rotation age (37 years) ^3^.

The major issue of curly birch propagation on plantations is that the first signs of ‘curliness’ among the trees appear at the age of 8 to 15 years ^6,7^. Until that time, it is necessary to maintain a large plantation of trees with unpredictable yield, since any stands of trees artificially planted using Karelian birch seeds always have an admixture of ordinary (non-curly) birches. Proportions of ‘curly’ trees among progenies of open-pollinated Karelian birch vary from 2–3% ^8^ to 25% (rarely 50%) ^9,10^. 60-70% of curly-wooded offspring appear from controlled crosses of two parents with a ‘curly’ phenotype ^11^.

Non-curly birches are usually removed from the stand at the age of about 10-13 years ^3^ to provide genuine curly trees with sufficient space and light. For this cleaning, a non-destructive visual assessment of curliness is commonly used ^12^. However, even at this age, tinning often encounters a problem, since detecting curliness by indirect morphological features of trees is rather difficult. Thus, understanding the genetic mechanism underlying the figured wood phenomenon of Karelian birch is becoming a truly challenging task, yet it has the potential to provide foresters with trait-specific molecular markers to help recognize curly birch genotypes at their earliest stages of development.

Throughout the long history of studying Karelian birch, various hypotheses have been suggested regarding the nature of the factors that determine curly wood ^12–14^. The heritability of the trait, though, is acknowledged by most researchers. Recently ^11^, have precisely evaluated the phenotypic segregation ratios in several progeny trials obtained from crosses of curly and non-curly birch parent trees. As a result, a simple Mendelian inheritance model for curliness was proposed: a monogenic trait with two alleles, the allele coding for curly phenotype is dominant over the allele coding for normal phenotype and the semi-dominant curly allele is lethal when homozygous ^11^.

With advances in modern high throughput genotyping technologies and well-developed experimental workflow of the Genome-Wide Association Study (GWAS), it is possible to take another step forward in the study of the molecular genetic basis of the curly birch phenotype. In the present study, we used 37,045 SNP markers to genotype 192 trees with and without the ‘curliness’ phenotype derived from Karelian birch crosses, to reveal how many loci could be potentially involved in ‘curliness’ formation and to search for genetic variants associated with this trait.

## 2. Materials and methods

### 2.1 Plant material and phenotyping

In the study we investigated the full-sib seed progeny of the Karelian birch, growing on the Zaonezhskaya Forest-Seed Plantation, Medvezhyegorsk region (62°15’N 35°02’E) in the south-eastern part of Karelia (Russia). 192 trees from two full-sib families derived from two controlled crosses between Karelian birch parents *№*133 × *№*134 and *№*51 × *№*58a have been analyzed (Supplementary Table S1). The crosses were carried out in 1987 and 2006, respectively, direct and reciprocal crosses were performed. “Curliness” (curly wood, cw) of phenotyped trees was determined by a nondestructive testing approach: by stem growth form and bark color, but mostly by the presence and shape of bulges on the stem surface. Among these, spherical thickened, finely tuberculate and also without signs of “curliness” are visually distinguished (Fig. 1). According to the type of the surface of the stem, one can roughly estimate the level of the manifestation of wood “curliness”. The most intense patterned texture in wood, as a rule, is formed in Karelian birch trees with a finely tuberculate type of stem surface. The signs of “curliness” in a pronounced form appear on average in the 8–10th year of the plant’s life. It was also established that as the plants develop, after 30–40 years the reverse process of “smoothing” or “swollen” of the previously convex stem surface is observed due to an increase in the thickness of the bark (Fig. 1c). Although these features make it difficult to reliably identify Karelian birch without wood destruction, tree breeding programs make extensive use of this non-destructive approach to detect “curliness” based on the externally visible stem morphology conducted on 10-year-old and older birches ^5,11^. The field studies including collection of plant material and phenotyping on birch plants presented in this paper comply with relevant institutional, national and international guidelines and legislation.

**Fig. 1.**
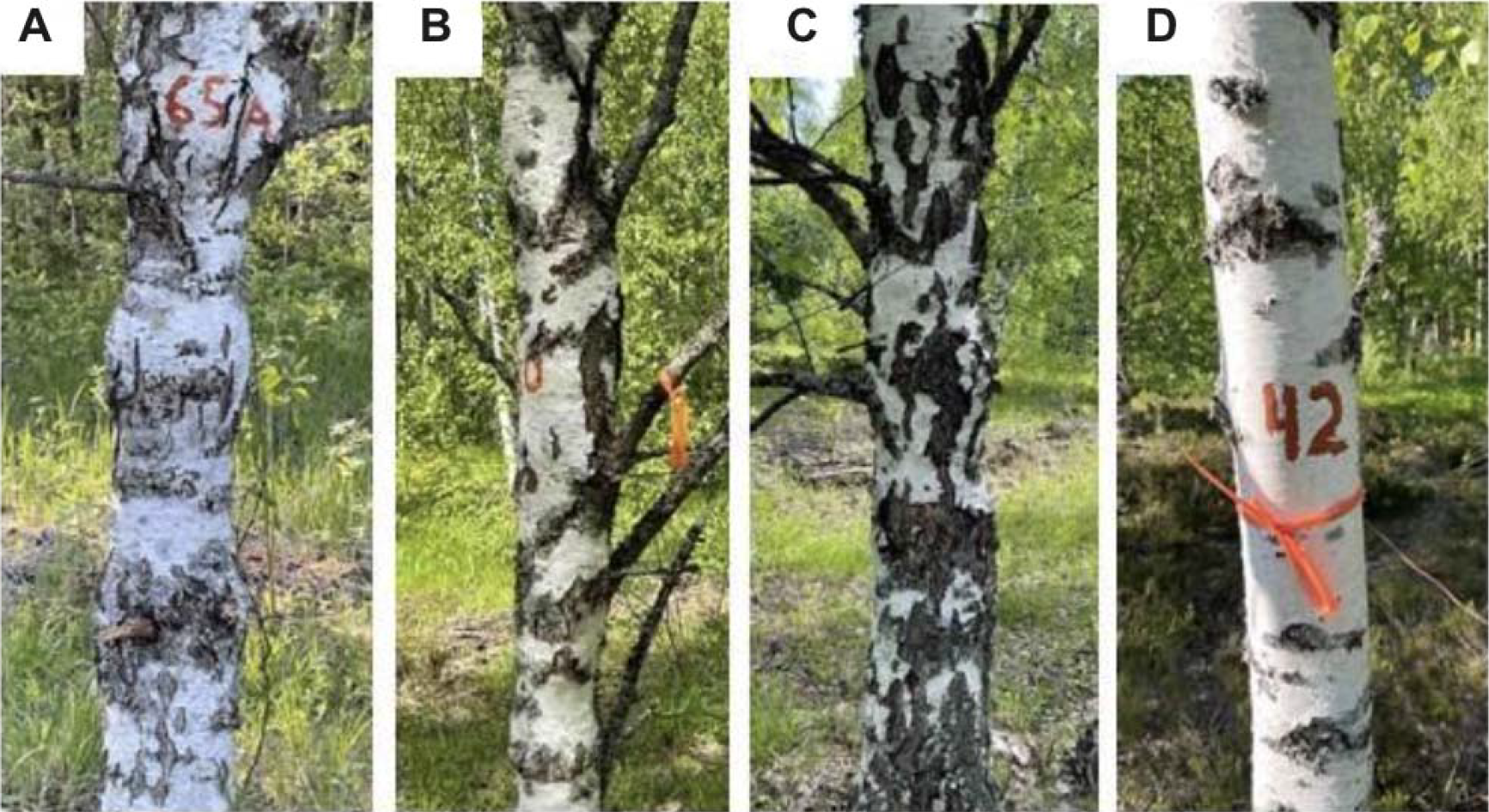
Types of the surface of the stem of the Karelian birch: spherical thickened (a), finely tuberculate (b), “swollen” (c) and without signs of “curliness” (d). Types of stem surface a, b and c were considered “curly” phenotype, d – as “non-curly” phenotype.

### 2.2 Genotyping Procedures

Genotyping-by-sequencing of plant material was performed using the RADseq method ^15^. A detailed protocol has been previously described and validated by our research group for the genotyping of various plant species, e.g. aspen ^16^, guar ^17^ and grapevine ^18^. Total DNA was isolated by a modified CTAB method according to the protocol ^19^ and stored at -20 °C until needed. The quality and concentration of the purified DNA samples were determined by gel electrophoresis and in the Qubit 3.0 Fluorometer (Thermo Fisher Scientific, Waltham, MA, USA). RADseq library was prepared with HindIII and NlaIII restriction enzymes for DNA digestion and sequenced using Illumina NovaSeq 6000 (Illumina, San Diego, CA, USA). Raw sequence data is available on NCBI SRA under the project number PRJNA997794. Tree numbers, corresponding barcodes and tree phenotypes are summarized in Supplementary Table S1.

### 2.3. Read Filtering, Alignment, and SNP Calling

The SNP calling procedure was performed using GATK v.4.4.0.0 software ^20^ after aligning reads on the reference genome assembly ^21^. Before alignment, the data was grouped and sorted using Picard version 3.0 (https://broadinstitute.github.io/picard/). Reads alignment was performed using a bowtie2 aligner with a very sensitive flag ^22^. Birch genome ^21^ sequence version 1.4c was used as a reference (https://genomevolution.org/coge/api/v1/genomes/35080/sequence). After alignment, SNP calling was performed with the GATK’s GenotypeGVCFs function using the parameter Max Alternate Alleles = 2. GATK’s Variant Filtration function was used to filter SNPs based on the following parameters: Minor Allele Frequency (removing SNPs with frequency > 0.05), Mapping Quality (removing SNPs with score < 40.0), Quality by Depth (removing SNPs with score/depth < 24.0), Fisher Strand Bias (removing SNPs with p-value > 60.0), Strand Odds Ratio (removing SNPs with odds ratio > 3.0), and Depth (removing SNPs with depth < 20.0).

### 2.4. Population Structure Assessment

In order to find the number of subpopulations within the studied birch cohort, ADMIXTURE software version 1.3 was used ^23^. The potential number of subpopulations was set from 1 to 15. The principal component analysis was performed using default parameters in the PLINK software version 1.9 ^24^. The number of principal components was set to 20. Principal components were estimated using the variance-standardized genetic relationship matrix. Linkage disequilibrium was evaluated using *r*^*2*^ which in turn was derived from the pairwise comparisons made in the PLINK software version 1.9. The *r*^*2*^ value was calculated for SNP pairs located within 1000 kb frames across the chromosomes. LOESS (locally estimated scatterplot smoothing) was used to estimate the regression between *r*^*2*^ and distance.

### 2.5. Association Mapping, SNP Annotation

To identify genotype-phenotype associations, the GAPIT version 3 software was used ^25^. The mapping was performed using the SUPER (Settlement of MLM Under Progressively Exclusive Relationship) approach ^26^. The first five principal components, as well as the kinship matrix calculated using methods described previously ^27^ were added to the model to account for the population structure and kin relationships, respectively. In order to assume an association, the adjusted p-value (Bonferroni correction) of less than 0.00000135 (0.05/37,045) was used, where 37,045 is the number of tests (SNPs). Additionally, false discovery rate (FDR) correction was used according to the Benjamini-Hochberg (BH) procedure. The SNPs that demonstrated BH-corrected p-values less than 0.05 were assumed to be associated with the phenotype. To scan for the candidate genes, the list of coding sequences was retrieved in the region on chromosome 10 restricted with SNPs S10_2623237 and S10_3848247 (+-30kb). Original annotation in gff format for assembled genome was used to extract potential candidates (https://genomevolution.org/coge/api/v1/downloads/?gid=35080&filename=Betula_pendula_subsp._pendula_annos1-cds0-id_typename-nu1-upa1-add_chr0.gid35080.gff). The proportion of the explained phenotype variance was estimated by performing an analysis of the variance of the linear model (lm() function) in the R base package.

### 2.6. SNP validation, molecular marker development

Sanger sequencing was applied to validate the high throughput SNP call performed by GATK using RADseq data. Four pairs of primers flanking the most significant SNPs were designed: for S10_3168885 forward 5′-GAAGAGAGGCTATAACGAGACATAAA and reverse 5′-CCTTCACTTACTAGCATGACTGAT; for S10_3472479 forward 5′-TCCTTCGGTCGATGAAATAACA and reverse 5′-CAATCTGCGAGAGGGAACAA; for S10_3465040 forward 5′-GGTTGGAAGAGCTCCATGAT and reverse 5′-GGAAGAATAAATAAGTCTGAGATGCC; for S10_3103637 forward 5′-GGGACAGTGCTGCATATCAT and reverse 5′-AAGGAAAGTCAATCTCACCAGAA. For the primer design *B. pendula* genome sequence version 1.4c ^21^ was used as a reference.

For all the designed primers PCR reaction mix contained 1 × Taq-buffer (2.5 mM Mg2+); 200 μM dNTP; 2.5 U Taq DNA polymerase (Eurogene, Moscow); 0.4 μM of each primer; ∼20 ng of template DNA, sterile distilled water in a final volume of 25 μl. PCR cycling conditions consisted of an initial denaturation step of 95 °C for 3 min, followed by 30 cycles of 95 °C for 30 s, 58 °C for 30 s, 72 °C for 45 s, and a final extension cycle at 72 °C for 5 min. PCR products were purified using magnetic particles, followed by termination PCR with BrilliantDye™ Terminator Cycle Sequencing Kit 3.1 using forward and reverse primer. Sanger sequencing was performed with Applied Biosystems 3500 Genetic Analyzer. The DNA sequences were aligned with UniproUGENE software ^28^. The genetic marker BpCW1 described in this paper is the subject of a patent application no. 2023122806 filed by the authors and protected by the Federal Service for Intellectual Property of Russia.

## 3. Results

### 3.1. Phenotyping, genotyping and SNP calling

The phenotypic evaluation was performed for 192 trees from two studied crosses. In the first full-sib population consisting of 88 trees, 70 plants were classified as ‘curly’ and 18 demonstrated non-curly phenotype. For the second cross (104 trees) 59 and 45 trees demonstrated curly and non-curly phenotypes, respectively (Supplementary Table S1).

The RADseq procedure yielded an average of 16,090,513 paired-end sequencing reads for each birch DNA sample. After the read filtering procedures, an average of 9,107,446 paired-end reads per plant were mapped to the birch genome. By analyzing alignment files, we identified 3,065,097 biallelic SNPs using the GATK SNP caller. As a result of filtering, 37,045 SNPs remained for further analysis of population structure and association mapping (Supplementary Table S2).

### 3.2. Population structure analysis revealed clusterization and rapid LD decay

To assess the population structure of the birch cohort we estimated the potential number of K clusters (sub-populations) using ADMIXTURE software ^23^. Next, we performed a principal component analysis implemented in PLINK software ^24^ to estimate the proportion of genotype variance explained by principal components as well as to perform visualization (Fig. 2A-D). Finally, we assessed a linkage disequilibrium decay using PLINK (Fig. 2E).

**Fig. 2.**
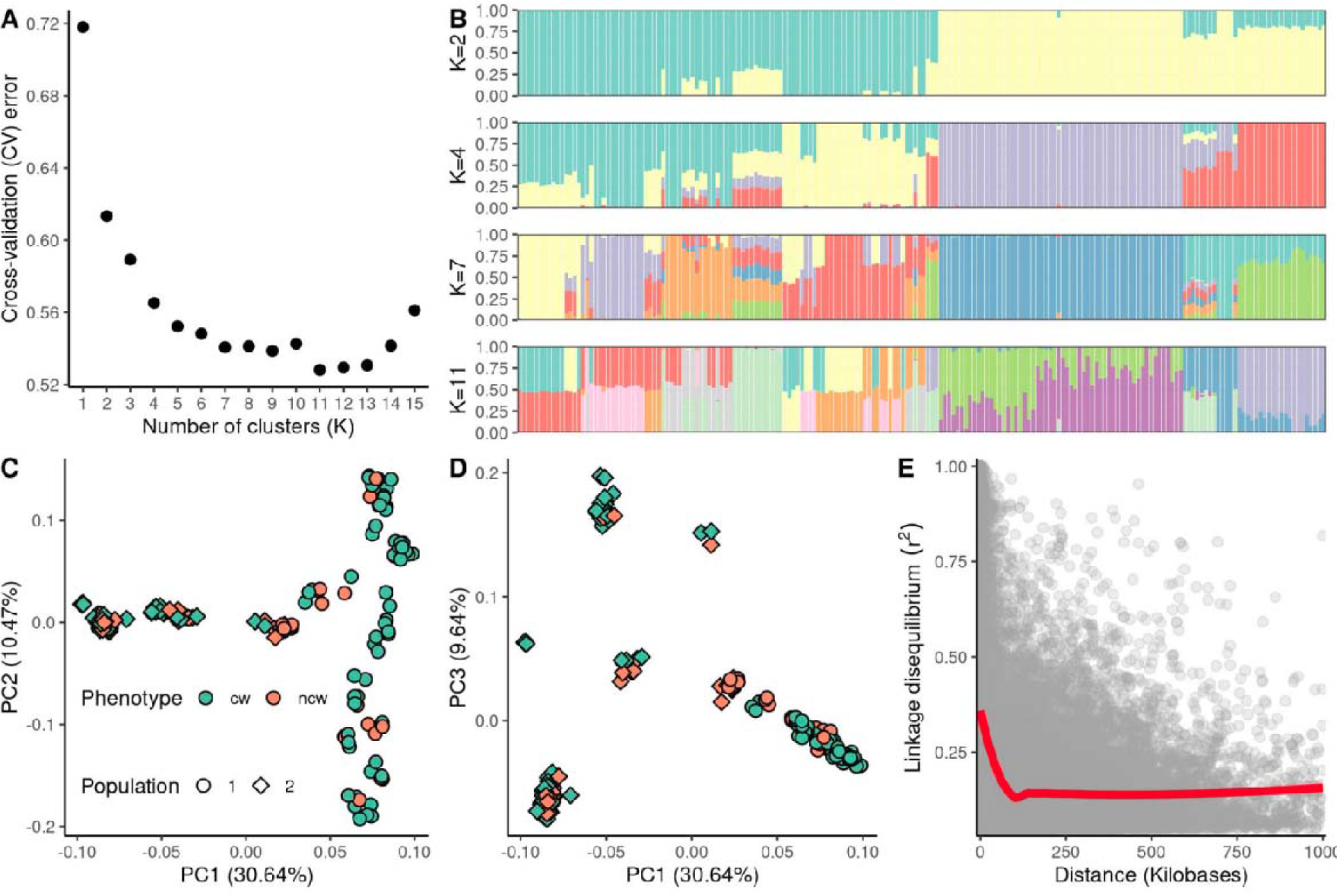
Population structure of the studied Karelian birch cohort. Panels A and B represent the results of the ADMIXTURE software. (A) Estimated cross-validation error value for possible clusters from 1 to 15. The drop in the cross-validation error indicates the optimal number of possible subpopulations. (B) ADMIXTURE bar plots reflecting the subpopulations at K = 2, 4, 7, 11 each bar corresponds to the birch accession, colors indicate the proportion of subpopulation admixtures in each accession. For K=2 Green color corresponds to the genetic admixtures from population 2, while yellow color corresponds to the genetic admixtures from population 1. Panels C and D represent the result of the Principal Component Analysis. Colors correspond to the phenotype of the trees (cw - ‘curly’, ncw - ‘non-curly’) and the shape indicates the corresponding full-sib population according to the plan of the birch plantation. The numbers in brackets correspond to the proportion of genotype variance explained by the principal component. (E) Linkage disequilibrium (LD) decay in the studied cohort of birch trees. Each dot corresponds to the *r*^*2*^ value between a pair of SNPs. The red line represents the LOESS curve. The grey marker corresponds to the 95% confidence interval.

The most significant drop of cross-validation (CV) error was observed when binning the birch cohort into two clusters (K=2) (Fig. 2A, B). This is in concordance with the information that the studied cohort was represented by two populations derived from two independent crosses. Subsequent binning at K=4, 7, 11 divides each of the two studied populations into different groups (Fig. 2B). The first three principal components explained 50.74% of genotypic variance which is in concordance with the strong population clustering (Fig. 2C, D). The first principal component explained the variation between the two studied populations obtained from two different crosses, while the second and the third principal components explained variation within them. Notably ‘curly’ and ‘non-curly’ phenotypes were evenly distributed between and within two full-sibs’ populations. Linkage disequilibrium (LD) decayed rapidly and was equal to 30.8 kb at *r*^*2*^ = 0.25 and 79.5 kb at *r*^*2*^ = 0.15 (Fig. 2E).

### 3.3. Association mapping revealed the target locus on chromosome 10

To perform association mapping, the SUPER model was applied accounting for population structure and relationships by introducing the first five principal components and kinship matrix. Out of the 37,045 SNPs tested, 3 SNPs (S10_3168885, S10_3465040, S10_3472479) were found to be significant after Bonferroni correction for multiple testing (Fig. 3).

**Fig. 3.**
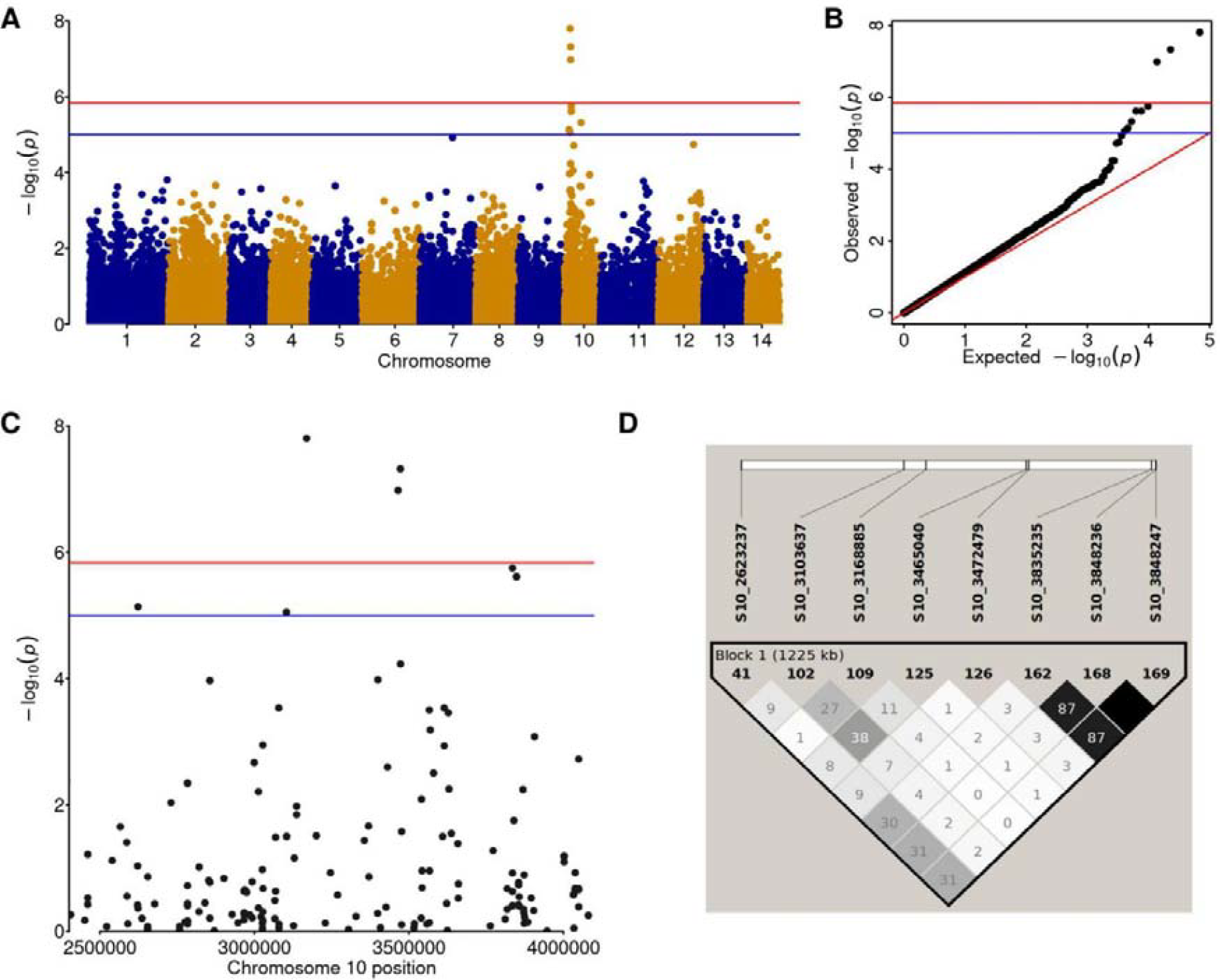
Results of the association mapping of curly wood trait. (A) Manhattan plot and respective (B) QQ-plot showing SNP markers associated with the ‘curly’ phenotype of Karelian birch. Each dot corresponds to a single SNP. Blue dots correspond to SNPs from odd chromosomes, yellow dots correspond to SNPs from even chromosomes. (C) Manhattan plot for the region of interest on chromosome 10 restricted with SNPs S10_2623237 and S10_3848247. The red line corresponds to the Bonferroni-adjusted significance threshold. The blue line corresponds to the FDR-adjusted significance threshold. Linkage between the associated SNPs (Panel D), numbers in squares correspond to the digits after the comma of the *r*^*2*^ value between a pair of SNPs.

Using a softer FDR threshold (α = 0.05) 5 additional SNPs (S10_2623237, S10_3103637, S10_3835235, S10_3848236, S10_3848247) surrounding the above-mentioned SNPs were identified to be significantly associated with the studied trait. 5 out of 8 SNPs were found within the coding sequences annotated for the *B. pendula* reference genome (Table 1). In addition, S10_3465040 was located in the 3’UTR region of the Bpev01.c0000.g0109 gene, at 27 bp downstream of the stop codon.

**Table 1.** SNPs associated with the ‘curly’ phenotype in two full-sib populations obtained from crossing Karelian birches according to GWAS results.

| SNP<br>chromosome_position | P.value | FDR | PVE,<br>% | Coding sequence,<br>putative function | Nearest upstream<br>transcript match | Nearest<br>downstream<br>transcript match |
| --- | --- | --- | --- | --- | --- | --- |
| S10_2623237 | 7.3E-06 | 0.0313 | 16.78 | Bpev01.c0000.g0035<br>unknown protein | Bpev01.c0000.g0034,<br>Alcohol<br>dehydrogenase | Bpev01.c0000.g0036,<br>RmlC-like cupins<br>superfamily protein |
| S10_3103637 | 8.85E-06 | 0.0337 | 37.25 | Bpev01.c0000.g0077,<br>3-ketoacyl-CoA<br>thiolase (exon) | Bpev01.c0000.g0076,<br>PB1 domain-<br>containing protein<br>tyrosine kinase | Bpev01.c0000.g0078,<br>mRNA splicing<br>factor thioredoxin-<br>like U5 snRNP |
| S10_3168885 | 1.57E-08 | 0.0005 | 25.24 |  | Bpev01.c0000.g0085,<br>Alba DNA/RNA-binding protein<br>LENGTH 405 | Bpev01.c0000.g0086,<br>O-Linked N-acetylglucosamine<br>transferase |
| S10_3465040 | 1.04E-07 | 0.0012 | 56.28 |  | Bpev01.c0000.g0109,<br>Unknown protein | Bpev01.c0000.g0110,<br>Pentatricopeptide<br>repeat-containing<br>protein |
| S10_3472479 | 4.77E-08 | 0.0008 | 22.74 |  | Bpev01.c0000.g0110,<br>Pentatricopeptide<br>repeat-containing<br>protein | Bpev01.c0000.g0111,<br>RING/U-box<br>superfamily protein<br>LENGTH 360 |
| S10_3835235 | 1.78E-06 | 0.0139 | 10.01 | Bpev01.c0000.g0140,<br>LanC-like protein 2 | Bpev01.c0000.g0139,<br>Plant protein of<br>unknown function<br>(DUF827) LENGTH<br>1345 | Bpev01.c0000.g0141,<br>SNF1-related protein<br>kinase |
| S10_3848236 | 2.42E-06 | 0.0139 | 9.03 | Bpev01.c0000.g0141,<br>SNF1-related protein<br>kinase | Bpev01.c0000.g0140,<br>LanC-like protein 2 | Bpev01.c0000.g0142,<br>Putative Peroxidase |
| S10_3848247 | 2.42E-06 | 0.0139 | 9.03 | Bpev01.c0000.g0141,<br>SNF1-related protein<br>kinase | Bpev01.c0000.g0140,<br>LanC-like protein 2 | Bpev01.c0000.g0142,<br>Putative Peroxidase |

At the next step we assessed the effect that the SNPs exerted on the phenotype. To do so the phenotype distribution was visualized (Fig. 4). For one of the most significant SNP (S10_3465040) the ‘non-curly’ phenotype was associated with ‘G’ allele in the homozygous state, while the presence of the alternative ‘A’ allele indicated the curly phenotype. Remarkably, this SNP showed the largest phenotypic variance explained (PVE=56.28%). That makes the S10_3465040 a good candidate for marker-assisted breeding applied to curly wood trait.

**Fig. 4.**
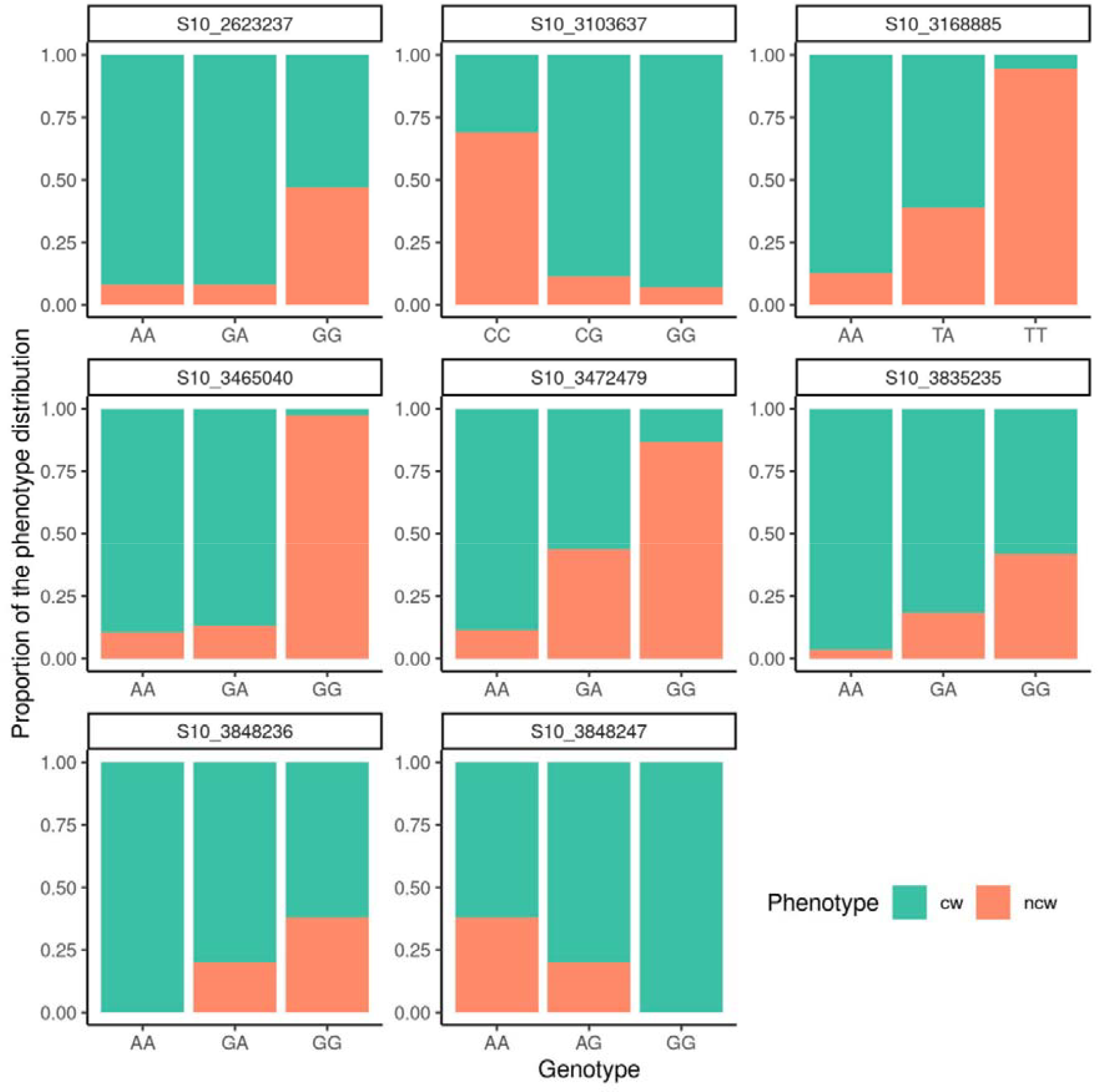
Barplots reflecting the effects of detected SNPs on the segregation of trees according to the “curly” phenotype. Each bar corresponds to a phenotype distribution across one of the three possible genotypes. Each colored bar shows the proportion of the phenotype distribution (cw - curly wood, ncw - non-curly wood) across genotypes.

Since all identified SNPs were located at the 10th chromosome, we assumed the presence of a locus associated with the studied trait (Fig. 3C). Despite the quite large chromosome region that carries SNPs associated with the trait (more than 1200 kb), the marginal SNPs (S10_2623237 and S10_3848247) demonstrated moderate (*r*^*2*^ > 0.3) linkage (Fig. 3D). Hence, we considered this region as potentially carrying a candidate gene or genes associated with the curly wood phenotype.

Detailed analysis of the region restricted by SNP S10_2623237 (3’-end) and SNP S10_3848247 (5’-end) plus an additional 30 kb at both ends, yielded 104 coding sequences (Supplementary Table S3). Most of the genes have been annotated as encoding unknown proteins, however, some genes from the list were previously discussed as potential players in curly wood formation.

For example, Bpev01.c0000.g0045 (Bpe_Chr10_2774789-2776265) encoding protein from Flavin-containing_monooxygenase_family located approx. 150 kb upstream from the candidate SNP S10_2623237. This coding sequence was recently reported as the *BpYucca5* gene involved in auxin biosynthesis ^29^. The *Yucca* gene family is a key player in IAA (the main natural auxin form) homeostasis. Much of IAA in plants is synthesized from tryptophan with the involvement of Yucca flavin-dependent monooxygenases ^30^. It has recently been proposed that the development of anomalous wood in Karelian birch may be associated with impaired auxin biosynthesis ^14,29^. Another possibly relevant gene Bpev01.c0000.g0081 encoding Sm-like_snRNP_protein (Bpe_Chr10_3118250-3127056) is located in 14613 bp downstream from SNP S10_3103637. Sm-like snRNP proteins are required for mRNA splicing, export, and degradation (He and Parker, 2000). It was shown that the homolog of the candidate gene Bpev01.c0000.g0081 in *Arabidopsis* (Sm-like protein (SAD1), AT5G48870) affects both ABA sensitivity and drought-induced ABA biosynthesis: ABA-deficiency was detected in sad1 mutant plants ^31^. The authors proposed a critical role for the SAD1 gene in RNA metabolism: sad1 mutation affects the decay rate of mRNA for an early component(s) in ABA signaling in drought stress conditions.

### 3.4. SNP validation revealed the first molecular marker for *B. pendula* curly wood trait (BpCW1)

PCR fragments obtained with primers flanking each of the four most significant SNPs were re-sequenced for the randomly chosen 16-20 birch trees. The accuracy of genotyping was estimated as 81%, 100%, 86%, 75% for SNPs S10_3103637, S10_3168885, S10_3465040, S10_3472479, respectively (Supplementary Table S4).

The length of PCR products obtained for neighboring SNPs S10_3465040 and S10_3472479 varied significantly in size in the populations due to numerous InDels detected within the amplified fragments. In particular, in the sequence of the PCR fragment containing S10_3472479, at least two InDels of 101 bp and 6 bp were detected by Sanger sequencing. PCR analysis of the entire sample of 192 trees with the primers specific for S10_3472479 revealed at least three more length variants of the PCR products (Supplementary Fig. S5). None of the InDels adjacent to S10_3472479 were significantly linked to the “curly” phenotype. As a consequence, no correlation between the PCR fragment size and the “curly” phenotype was detected (Supplementary Fig. S5).

At least 3 InDels of 54 bp, 13 bp and 2 bp were detected within the PCR products amplified with primers flanking S10_3465040. Notably, the 54 bp deletion both in the homozygous and heterozygous state correlated perfectly with the “curly” phenotype in the analyzed populations. 54 bp deletion results in a 476 bp amplified fragment compared to the size of 530 bp intact PCR fragment, which makes it easily recognizable by electrophoresis (Fig. 5). Of the 190 trees tested for the presence of a 54 bp deletion with this PCR marker, 174 trees (92%) were identified correctly as ‘curly’ or ‘non-curly’ depending on the presence or absence of the 476 bp PCR-fragment (Supplementary Fig. S6). To our knowledge, this is the first molecular marker proposed for the recognition of *Betula pendula* Roth var. *carelica* with the curly wood phenotype (BpCW1).

**Fig. 5.**
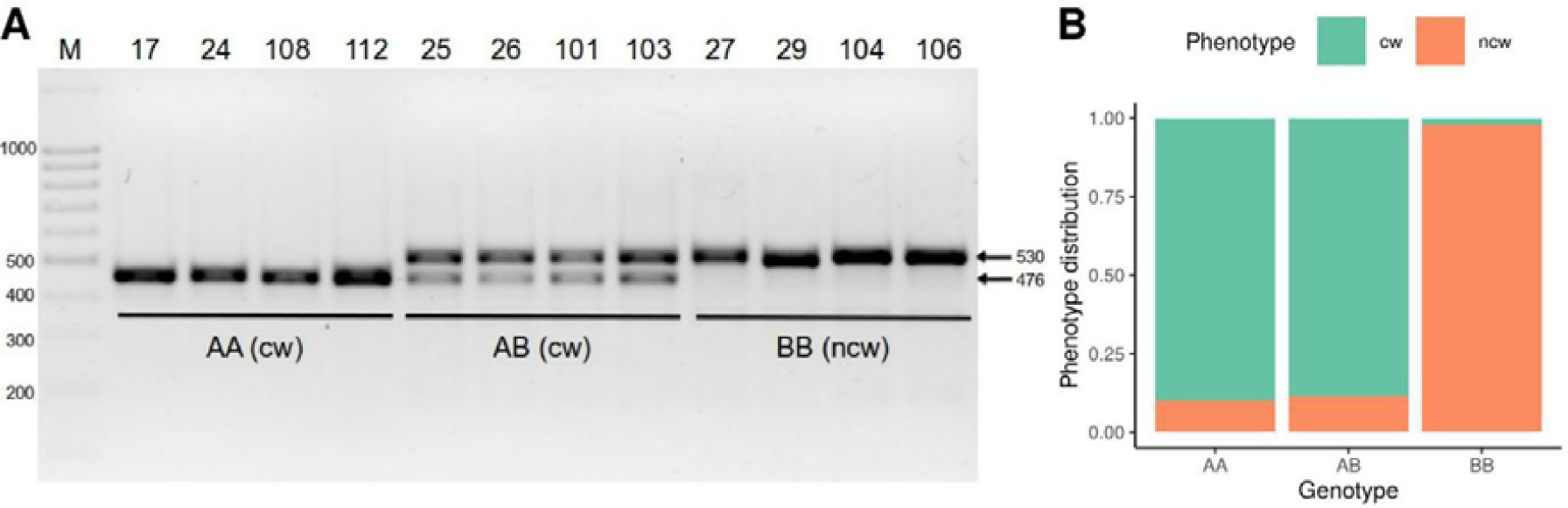
Results of the validation of the 54 bp deletion associated with SNP marker S10_3465040. (A) The result of PCR amplification of DNA fragment containing deletion associated with the curly wood phenotype. Samples 17, 24, 108, 112 and 25, 26, 101, 103 demonstrating curly wood (cw) phenotype carry deletion in homozygous (AA) and heterozygous state (AB), respectively. Samples 27, 29, 104, 106 do not carry deletion (BB) and demonstrate non-curly wood phenotype (ncw). (B) Phenotype (cw, ncw) distribution among genotypes associated with the deletion (AA, AB, BB) for the studied cohort of 192 trees.

## Discussion

In this study we attempted to unravel the enigma of genetic factors underlying the phenomenon of curly wood in Karelian birch, which has been debated for over a century ^32^. Although in 1922 it was already shown that the patterned texture of the wood is quite stably transmitted to progenies during seed propagation ^33^, there were many adherents of the ‘infectious’ or ‘pathological’ hypotheses, which considered curly wood to be the result of exposure to biotic or abiotic stressors ^34–36^. The idea finds support in some modern studies, for example, the influence of a low-temperature stress gradient is assumed, which determines the boundaries of the distribution of Karelian birch ^37^. According to the reports, the suppression of xylem-phloem transport and a decrease in the rate of plant growth occur in parallel with an increasing deficit of heat, moisture, or mineral nutrition. Recently, an ecological and genetic hypothesis has been proposed, according to which the origin of the Karelian birch is associated both with the natural and climatic conditions of its growth and with genetic factors ^38,39^.

T. Ruden (1954) was the first to postulate that the ‘curly’ trait is monogenic, the gene can be dominant and lethal when homozygous ^40^. Later it was proposed that the inheritance of curly wood is due to a series of multiple alleles in one locus or genes in different loci ^41^. In 2011 a hypothesis about the epigenetic origin of curly wood of the Karelian birch was suggested ^42^.

Since the aberrant ‘curly’ phenotype occurs regularly, albeit sporadically within the north-western part of the silver birch distribution area, and is inherited in offspring, the hypothesis that curly birch can be a single-locus mutant of *B. pendula* seems quite plausible. This hypothetical mutation apparently disrupts the normal development of the plant, affecting not only the general habitus but probably also the tree’s longevity. As stated by some reports, the life cycle of the Karelian birch takes approximately 50–60 years ^35,39,43,44^, while the life cycle of the silver birch takes 120–140 years. The impairment of xylogenesis observed in Karelian birch may be due to the loss of function of a particular causative gene. It may also play a role in whether this mutation, which affects the fitness of the tree, is present in a homo- or heterozygous state. Accordingly, Kärkkäinen et al. (2017) reported that their analysis of crosses studying the inheritance of curly birch phenotype indicates a Mendelian inheritance based on a single semi-dominant gene that is lethal when homozygous ^11^.

Consistent with this report, our study based on RADseq and association mapping revealed a single interval on chromosome 10 spanning ∼1200 kb, where 8 SNPs were found significantly associated with the trait under study. The fact that one of the SNP markers (S10_3465040) was identified explaining up to 56% of phenotypic variation (Table 1) speaks in favor of the previously suggested monogenic (Mendelian) mode of inheritance for the curly wood trait. This particular SNP S10_3465040 and the 7 kb distant SNP S10_3472479 seem to fall into a region on the chromosome rich in small InDels. Three InDels were found only within 530 bp surrounding SNP S10_3465040, and even more within the 621 bp interval surrounding SNP S10_3472479.

Thus, in the region of interest, we observed a distinct structural characteristic: a high frequency of insertions and deletions of varying sizes. This structural dynamism might hint at the active presence of transposable elements (TEs) within this region. TEs, often termed “jumping genes”, have been recognized for their capacity to move within the genome, and their activation can be influenced by various factors. TEs are not just passive genomic elements; they can play pivotal roles in gene regulation, especially in response to environmental stresses ^45^. Furthermore, TEs can influence alternative splicing patterns, potentially serving as alternative splicing start or acceptor sites ^46^.

We found that the 54 bp InDel located at a distance of 160 bp from SNP S10_3465040 successfully predicted the “curly” phenotype for 92% of the 190 trees examined. It should be considered that the phenotyping of the “curly” trait was carried out by the method of non-destructive visual assessment of the habitus of the tree, without assessing the cut, so a certain percentage of erroneous phenotyping can be expected. Notably, of the 16 mislabeled trees, 15 were diagnosed as “curly” by the PCR marker, while visual assessment phenotyped these trees as “non-curly”. Only one tree was mislabeled the other way around. This 54 bp InDel, however, does not fall into any coding sequence, although it is located in the 3’UTR region of the Bpev01.c0000.g0109 gene encoding an unknown protein. Blast analysis of the Bpev01.c0000.g0109 sequence revealed just a few hits – in the genomic assemblies of *Quercus, Alnus* and *Fagus*, which suggests that it may be specific to the *Fagales* order. Deletions in the 3’UTR region of this unknown gene can dramatically affect the fate of its mRNAs. The 3′-UTR regulatory regions are involved in polyadenylation, stability, transport and mRNA translation in plant cells ^47^. When the effect of increasing the length of the 3’UTR on expression of luciferase mRNA was examined in transiently transfected carrot protoplasts, expression increased 18-fold when the length of the 3’-UTR was increased from 7 to 27 bases ^48^.

Several studies have highlighted the profound effects of TE insertions, especially within the 3’UTR regions of genes. Such insertions can lead to translational repression, thereby influencing phenotypic outcomes. For instance, in rice, the Ghd2 gene, a regulator of flowering time, is impacted by such a mechanism ^49^. This phenomenon isn’t exclusive to plants. In Drosophila, a TE insertion has been linked to enhanced survival rates in response to chemicals, due to its effect on the upregulation of the CG11699 gene ^50^. Similarly, in mammals, TEs, particularly SINE repeats, have been shown to modulate gene promoter activity ^51^. A study in humans demonstrated that the expression of an AcGFP reporter gene is significantly attenuated by inverted Alu repeats in the 3′UTR ^52^. Given the proximity of this SNP to the 3’UTR and the known effects of TEs on gene regulation, it can be assumed that TEs may influence the curly wood phenotype of the Karelian birch.

Bpev01.c0000.g0109 can be considered as one of the possible candidate genes, however, linkage disequilibrium (LD) between the S10_3465040 and the putative causal locus located near the identified polymorphism can also be assumed. Rapid LD decay was reported for many arboreal species including pines ^53^, aspens ^54^ and willows ^55^ due to the high level of genetic diversity (heterozygosity) of these woody species. In our study, though, the identified LD at *r*^*2*^ = 0.15 was higher (79.5kb) compared to one revealed in the previous study (23kb) performed on the 60 genetically diverse birch trees ^21^. This could be explained by the fact that the association study was performed using a cohort of related individuals (two populations of full-sibs), which undergo only few recombination events. This may be the reason why the chromosomal interval at LG10, where significantly associated SNPs were detected, stretched to 1200 kb.

The detected interval on chromosome 10 was discussed previously as containing the sequence of the *BpYucca5* gene involved in auxin biosynthesis which is critical for normal xylogenesis in Karelian birch ^29^. The *BpYucca5* gene is listed as one of the 104 coding sequences detected within the interval on chromosome 10. Among the potential candidates, there are also several genes encoding proteins that facilitate the processing, splicing, editing, stability and translation of mRNAs (Supplementary Table 3). Some of them are regulated by abiotic stress factors, such as Sm-like_snRNP_protein (Bpe_Chr10_3118250-3127056) which affects ABA biosynthesis under drought conditions.

Assuming that a causal polymorphism (in which TEs may be involved) can influence gene expression or mRNA stability depending on the environment, it may be explained why the Mendelian inheritance of this valuable trait has long been contested. Indeed, the formation of figured patterns in the wood of Karelian birch can begin at any age: cuts of trees are described in which patterned wood began to form at 8, 15, and even at 25 years. Moreover, there are examples of cross-cuts of trees, where the formation of a patterned wood texture is visible only in a certain sector of the trunk, perhaps, due to the special light conditions for the growth of the tree ^2^.

The phenomenon, when the degree of a trait expression differs in individuals of the same genotype, has been described as “expressivity” ^56^. Thus, the dominant allele may underlie the cause of figured wood, however, the degree of xylogenesis anomalies and the stage of development at which they occur depend on environmental conditions. That may explain the reason why the segregation ratio of curly and non-curly progenies in the crossing of Karelian birches varies from study to study ^2,57^.

The exact causes of expressivity are still not well understood, although they most likely lie in the molecular mechanisms governing genetic regulation ^56^. Our results suggest that the variation of the “curly” phenotype in Karelian birch is linked to the single interval on chromosome 10, where a high frequency of insertions and deletions of different sizes was observed. Unfortunately, the RADseq sequencing approach employed in our study doesn’t provide a comprehensive resolution of this region. To determine the true causal polymorphism in this interval, a deeper resequencing effort is required. However, the discovered PCR marker BpCW1 can even now be recommended for selection at an early stage of development of the Karelian birch seedlings, which later, upon reaching the age of 8-10 years, will most likely show the characteristic features of decorative (patterned) wood.

Identification of genetic determinants of curly wood phenotype and development of corresponding molecular markers can prevent excessive costs and efforts in the maintenance of Karelian birch plantations for industrial purposes and, possibly, reduce the anthropogenic pressure on the natural populations of these trees.

## Acknowledgments

We thank Dr. Anna Igolkina for the valuable discussion about the role of transposable elements in gene expression and trait manifestation.

## Ethics approval and consent to participate

All methods in the present research were performed in accordance with the relevant guidelines and regulations specified in Nature Portfolio journals’ Editorial policies. The present research did not involve human or animal participants.

## Research involving plants, permission for collection of plant material

The research and field studies on plants presented in this paper comply with relevant institutional, national and international guidelines and legislation. Karelian birch (*Betula pendula* Roth var. *carelica* (Mercklin) Hämet-Ahti) is a protected birch variety in Republic of Karelia, permission for the collection of the plant specimens was issued by the Ministry of Natural Resources and Environment of the Republic of Karelia.

## Data availability

The high-throughtput sequencing datasets generated during the current study are available in the NCBI’s Sequence Read Archive repository, https://www.ncbi.nlm.nih.gov/bioproject/PRJNA997794.

## Bibliography

1. Hagqvist, R. & Mikkola, A. Visakoivun kasvatus ja käyttö. (Metsäkustannus & Visaseurary: Hämeenlinna, 2008).

2. Vetchinnikova, L. V. & Titov, A. F. The Karelian Birch: a Unique Biological Object. Biol. Bull. Rev. 10, 102–114 (2020).

3. Viherä-Aarnio, A. & Hagqvist, R. Curly birch (betula pendula var. carelica), wooden “marble” from finland-soon easily available. in Proceedings of the international scientific conference on hardwood processing (2017).

4. Hynynen, J. et al. Silviculture of birch (Betula pendula Roth and Betula pubescens Ehrh.) in northern Europe. Forestry 83, 103–119 (2010).

5. Vetchinnikova, L. V., Titov, A. F. & Kuznetsova, T. Yu. KarelLJskaiLJaLJ bereza: biologicheskie osobennosti, dinamika resursov i vosproizvodstvo = Curly birch: biological characteristics, resource dynamics, and reproduction. (KarellJskiĭ nauchnyĭ tlJslJentr RAN, 2013).

6. Ermakov, V. Mekhanizmy adaptatsii berezy k usloviyam Severa[Mechanisms of birch adaptation to the conditions of the North]. (Izd-vo” Nauka,” Leningradskoe otd-nie, 1986).

7. Ermakov, V. & Vetchinnikova, L. Growing birch wood with combined grain. Drevarsky Vysk. 46, 37–42 (2001).

8. Johnsson, H. Avkommor av Masur bjork (Experiments with Masur Birch). Sartryck Ur Sven. Skogsvardsforeningens Tidskr. 12 (1951).

9. Heikinheimo, O. Experiences in the growing of curly birch. Commun Inst Fenn 39, 1– 26 (1951).

10. Larsen, C. Curly Birch. Dan. Skovforen. Tidsskr. 33–72 (1940).

11. Kärkkäinen, K., Viherä-Aarnio, A., Vakkari, P., Hagqvist, R. & Nieminen, K. Simple inheritance of a complex trait: figured wood in curly birch is caused by one semidominant and lethal Mendelian factor? Can. J. For. Res. 47, 991–995 (2017).

12. Vetchinnikova, L. V. & Titov, A. F. Curly Birch: Some Secrets Remain. Biol. Bull. Rev. 13, 162–174 (2023).

13. Novitskaya, L. L. Regeneration of bark and formation of abnormal birch wood. Trees 13, 74–79 (1998).

14. Novitskaya, L. L. et al. The Formation of Structural Abnormalities in Karelian Birch Wood is Associated with Auxin Inactivation and Disrupted Basipetal Auxin Transport. J. Plant Growth Regul. 39, 378–394 (2020).

15. Peterson, B. K., Weber, J. N., Kay, E. H., Fisher, H. S. & Hoekstra, H. E. Double digest RADseq: an inexpensive method for de novo SNP discovery and genotyping in model and non-model species. PloS One 7, e37135 (2012).

16. Zhigunov, A. V. et al. Development of F1 hybrid population and the high-density linkage map for European aspen (Populus tremula L.) using RADseq technology. BMC Plant Biol. 17, 180 (2017).

17. Grigoreva, E. et al. Development of SNP Set for the Marker-Assisted Selection of Guar (Cyamopsis tetragonoloba (L.) Taub.) Based on a Custom Reference Genome Assembly. Plants 10, 2063 (2021).

18. Volynkin, V. et al. The Assessment of Agrobiological and Disease Resistance Traits of Grapevine Hybrid Populations (Vitis vinifera L. × Muscadinia rotundifolia Michx.) in the Climatic Conditions of Crimea. Plants 10, 1215 (2021).

19. Rahimah, A. R., Cheah, S. C. & Rajinder, S. Freeze drying of oil palm (Elaeis guineensis) leaf and its effect on the quality of extractable DNA. J. Oil Palm Res. 18, 296–304 (2006).

20. McKenna, A. et al. The Genome Analysis Toolkit: a MapReduce framework for analyzing next-generation DNA sequencing data. Genome Res. 20, 1297–1303 (2010).

21. Salojärvi, J. et al. Genome sequencing and population genomic analyses provide insights into the adaptive landscape of silver birch. Nat. Genet. 49, 904–912 (2017).

22. Langmead, B. & Salzberg, S. L. Fast gapped-read alignment with Bowtie 2. Nat. Methods 9, 357–359 (2012).

23. Alexander, D. H., Novembre, J. & Lange, K. Fast model-based estimation of ancestry in unrelated individuals. Genome Res. 19, 1655–1664 (2009).

24. Purcell, S. et al. PLINK: a tool set for whole-genome association and populationbased linkage analyses. Am. J. Hum. Genet. 81, 559–575 (2007).

25. Lipka, A. E. et al. GAPIT: genome association and prediction integrated tool. Bioinformatics 28, 2397–2399 (2012).

26. Wang, Q., Tian, F., Pan, Y., Buckler, E. S. & Zhang, Z. A SUPER Powerful Method for Genome Wide Association Study. PLoS ONE 9, e107684 (2014).

27. VanRaden, P. M. et al. Invited review: reliability of genomic predictions for North American Holstein bulls. J. Dairy Sci. 92, 16–24 (2009).

28. Okonechnikov, K., Golosova, O., Fursov, M., & UGENE team. Unipro UGENE: a unified bioinformatics toolkit. Bioinforma. Oxf. Engl. 28, 1166–1167 (2012).

29. Tarelkina, T. V. et al. Expression Analysis of Key Auxin Biosynthesis, Transport, and Metabolism Genes of Betula pendula with Special Emphasis on Figured Wood Formation in Karelian Birch. Plants 9, 1406 (2020).

30. Ljung, K. Auxin metabolism and homeostasis during plant development. Development 140, 943–950 (2013).

31. Xiong, L. & Zhu, J.-K. Regulation of Abscisic Acid Biosynthesis. Plant Physiol. 133, 29–36 (2003).

32. Vetchinnikova, L. V. & Titov, A. F. Genesis of the karelian birch. an ecogenetic hypothesis. Ecol. Genet. 14, 3–18 (2016).

33. Hintikka, T. J. Die,, Wisa”-Krankheit der Birken in Finnland. Z. Für Pflanzenkrankh. Gallenkd. 32, 193–210 (1922).

34. Atanasoff, D. Virus stem pitting of birch (Wisa and Maser Disease). Z. Für Pflanzenkrankh. Pflanzenpathol. Pflanzenschutz 74, 205–208 (1967).

35. Saks, K. & Bander, V. New data about the origin of Karelian birch. Tr Inst Ekol Rast Zhivotn (1975).

36. Vailionis, L. Lietuvos berzu reta. Referat: Die Wisakrankheit in den Wäldern Litauens. Kaunas Sr Hort Bot Univ 3, 5–36 (1935).

37. Vetchinnikova, L. V. & Titov, A. F. The origin of the Karelian birch: An ecogenetic hypothesis. Russ. J. Genet. Appl. Res. 7, 665–677 (2017).

38. Vetchinnikova, L., Kharin, V., Spektor, E. & Bumagina, Z. MULTIVARIATE ANALYSIS OF INHERITING FEATURES OF PATTERNED WOOD TEXTURE IN HYBRID PROGENY OF CURLY (KARELIAN) BIRCH. Russ. J. For. Sci. 4, 70–74 (2003).

39. Vetchinnikova, L. V. KarelLJskaiLJaLJ bereza i drugie redkie predstaviteli roda Betula L. (Nauka, 2005).

40. Ruden, T. On speckled birch (“mazer-birch”) and some other forms of curled birch. Meddelelser Fra Det Nor. Skogforsøksves. 43, 455–505 (1954).

41. Johnsson, H. Genetic characteristics of Betula verrucosa Ehrh. and B. pubescens Ehrh. Ann. For. 6, 91–133 (1974).

42. Mashkina, O. S., Butorina, A. K. & Tabatskaya, T. M. Karelian birch (Betula pendula Roth. var. carelica Merkl.) as a model for studying genetic and epigenetic variation related to the formation of patterned wood. Russ. J. Genet. 47, 951–957 (2011).

43. Ermakov, V. Reproduction of Karelian birch by inoculation method. Lesn. Genet. Sel. Semenovod. For. Genet. Sel. Seed Breed. 282–293 (1970).

44. Mikkelä, H. Guide to the Montell trail in the Punkaharju experimental area. (Metsäntutkimuslaitos, 1996).

45. Reddy, A. S. N., Marquez, Y., Kalyna, M. & Barta, A. Complexity of the Alternative Splicing Landscape in Plants. Plant Cell 25, 3657–3683 (2013).

46. Casacuberta, E. & González, J. The impact of transposable elements in environmental adaptation. Mol. Ecol. 22, 1503–1517 (2013).

47. Bernardes, W. S. & Menossi, M. Plant 3’ Regulatory Regions From mRNA-Encoding Genes and Their Uses to Modulate Expression. Front. Plant Sci. 11, 1252 (2020).

48. Tanguay, R. L. & Gallie, D. R. The effect of the length of the 3′-untranslated region on expression in plants. FEBS Lett. 394, 285–288 (1996).

49. Shen, J. et al. Translational repression by a miniature inverted-repeat transposable element in the 3′ untranslated region. Nat. Commun. 8, 14651 (2017).

50. Mateo, L., Ullastres, A. & González, J. A Transposable Element Insertion Confers Xenobiotic Resistance in Drosophila. PLoS Genet. 10, e1004560 (2014).

51. Estécio, M. R. H. et al. SINE Retrotransposons Cause Epigenetic Reprogramming of Adjacent Gene Promoters. Mol. Cancer Res. 10, 1332–1342 (2012).

52. Fitzpatrick, T. & Huang, S. 3’-UTR-located inverted Alu repeats facilitate mRNA translational repression and stress granule accumulation. Nucleus 3, 359–369 (2012).

53. Brown, G. R., Gill, G. P., Kuntz, R. J., Langley, C. H. & Neale, D. B. Nucleotide diversity and linkage disequilibrium in loblolly pine. Proc. Natl. Acad. Sci. 101, 15255– 15260 (2004).

54. Ingvarsson, P. K. Nucleotide Polymorphism and Linkage Disequilibrium Within and Among Natural Populations of European Aspen (Populus tremula L., Salicaceae). Genetics 169, 945–953 (2005).

55. Berlin, S., Fogelqvist, J., Lascoux, M., Lagercrantz, U. & Rönnberg-Wästljung, A. C. Polymorphism and Divergence in Two Willow Species, Salix viminalis L. and Salix schwerinii E. Wolf. G3 GenesGenomesGenetics 1, 387–400 (2011).

56. Miko, I. Phenotype variability: penetrance and expressivity. Nat. Educ. 1, 137 (2008).

57. Korovin, V., Novitskaya, L. & Kurnosov, G. Structural abnormalities of the stem in woody plants. Mosc. State For. Univ. Mosc. 258, (2003).

